# Characterization of Heat Responsive microRNAs and Phased Small Interfering RNAs in Reproductive Development of Flax

**DOI:** 10.1101/2021.10.02.461750

**Authors:** Suresh Pokhrel, Blake C. Meyers

**Affiliations:** Division of Plant Sciences, University of Missouri, Columbia, MO 65211; Donald Danforth Plant Science Center, St. Louis, MO 63132

## Abstract

Plants will face increased heat stress due to rising global temperatures. Heat stress affects plant reproductive development and decreases productivity; however, the underlying molecular mechanisms of these processes are poorly characterized. Plant small RNAs (sRNAs) have important regulatory roles in plant reproductive development following abiotic stress responses. We generated sRNA transcriptomes of three reproductive bud stages at three different time points to identify sRNA-mediated pathways responsive to heat stress in flax. With added sRNA transcriptomes of vegetative tissues, we comprehensively annotated miRNA and phasiRNA-encoding genes (*PHAS*) in flax. We identified 173 miRNA genes, of which 42 are novel. Our analysis revealed that 141 miRNA genes were differentially expressed between tissue types while 18 miRNA genes were differentially expressed in reproductive tissues following heat stress, including members of miR2118/482 and miR2275 families, known triggers of reproductive phasiRNAs. Furthermore, we identified 68 *21-PHAS* flax loci from protein coding and non-coding regions, four *24-PHAS* loci triggered by miR2275, and 658 24-*PHAS*-like loci with unknown triggers, derived mostly from non-coding regions. The reproductive phasiRNAs are mostly downregulated in response to heat stress. Overall, we found that several previously unreported miRNAs and phasiRNAs are responsive to heat stress in flax reproductive tissues.

## Introduction

Agricultural production is negatively impacted by climate change, especially because of higher temperatures. The global yield of three main staple crops is estimated to be reduced by 3.2 to 6.0% per 1°C increase in global mean temperature (Zhao et al., 2017). The effect of higher temperature is greater on reproductive development, especially through impacts on pollen development, pollination, and embryo formation (Hedhly et al., 2009). Although there are studies that assess the impacts of higher temperature on growth, physiological processes, and yields on crops such as wheat, maize, rice, canola, and flax (Cross et al., 2003; Wang et al., 2016; Zhao et al., 2017; Pokharel et al., 2020), limited knowledge is available on the underlying molecular mechanisms that play roles in heat stress; such knowledge would enable the development of thermo-tolerant crops.

Heat shock proteins (HSPs) are major players in heat tolerance (Hasanuzzaman et al., 2013). HSPs are produced in response to heat stress after activation by heat stress transcription factors (HSFs) such as MULTIPROTEIN BRIDGING FACTOR1C, WRKYGQK motif–containing proteins, and Basic leucine zipper (Zhang et al., 2017; He et al., 2019). Multiple thermosensors initiate HSF activation after sensing heat stress. The conserved heat tolerance process in plants involves regulatory cascades of heat-responsive genes as well as other regulators such as non-coding RNAs that play a role in modulation of gene expression and these non-coding molecules are yet to be identified.

Plant sRNAs are implicated in modulating expression of genes related to heat stress (He et al., 2019). These sRNAs are regulatory elements that modulate gene expression, genome-wide, by transcriptional and post-transcriptional gene silencing. Typically, sRNAs are 20 to 24 nucleotides (nt) in length and are diced by DICER-LIKE proteins (DCLs) from hairpin or double stranded RNA precursors formed by RNA-DEPENDENT RNA POLYMERASEs (RDRs). The sRNAs form a silencing complex with Argonaute (AGO) proteins to modulate the expression of genes and transposable elements (TEs). The major classes of plant sRNAs are: microRNAs (miRNAs), phased secondary small interfering RNAs (phasiRNAs) and secondary small interfering RNA (siRNAs) (Fei et al., 2013).

The miRNAs are the master regulators to attenuate gene expression by transcript cleavage and degradation or translation inhibition in a homology-dependent manner (Zhai et al., 2011). There have been reports of heat responsive miRNAs such as miR398, miR156, miR159, miR160, miR167, miR396, and miR408 during vegetative and reproductive stages of plants like Arabidopsis, sunflower, maize, and tomato (Guan et al., 2013; Stief et al., 2014; Lin et al., 2018; Giacomelli et al., 2012; He et al., 2019; Keller et al., 2020). phasiRNAs are generated from mRNA transcripts of coding and non-coding regions of the genome mainly after the cleavage of trigger miRNAs. A special class of phasiRNAs that are produced from transacting siRNA loci (TAS) are called tasiRNAs (Fei et al., 2013). In Arabidopsis, tasiRNAs from the TAS1 locus are found to target HEAT-INDUCED TAS1 TARGET1 (HTT1) and HTT2 proteins that accumulate in response to heat stress (Li et al., 2014). Complete loss of reproductive 24-nt phasiRNAs after knock out of *DCL5*, a reproductive phasiRNA pathway gene, in maize, produces male sterility at high temperatures (Teng et al., 2020) while mutation in reproductive 21-nt phasiRNAs-generating loci produces photoperiod- and thermo-sensitive male sterility in rice (Fan et al., 2016; Zhou et al., 2012).

Flax is a widely cultivated crop species from the family Linaceae known for oil, fiber, and medicinally important compounds. Heat stress in flax reduces boll formation and seed set, and it may be due to the failure of pollen tubes to reach ovules (Cross et al., 2003). In this study, flax plants were subjected to 7 day and 14 day heat treatments after flower initiation in a growth chamber to simulate heat stress conditions in the field. We harvested three reproductive bud stage tissues at three time points and generated sRNA transcriptomes to identify sRNA-mediated pathways responsive to heat stress. With added sRNA transcriptomes of leaves, stem, and root, we comprehensively annotated miRNA- and phasiRNA-spawning genes (*PHAS*) in flax. We found that several miRNAs and both types of reproductive phasiRNAs (21 nt and 24 nt) are mostly down-regulated in response to heat stress. Overall, we found that several previously unreported flax miRNAs and phasiRNAs are responsive to heat stress in reproductive tissues.

## Results

### Comprehensive annotation of miRNA in flax

In the latest release of miRBase version 22 (http://www.mirbase.org), there are 124 miRNA precursors deposited for flax, mainly identified from computational approaches (Barvkar et al., 2013). There are studies which annotated flax miRNAs based on sRNA sequencing in control, nutrient, saline, and alkaline stress conditions (Yu et al., 2016; Melnikova et al., 2016) but their miRNA lists aren’t available from miRBase; they predicted a high number of novel candidate miRNAs (233 to 475), in addition to observing known miRNAs, suggesting that most of them may be false positives. Since the highly-studied model plant Arabidopsis only has a few hundred miRNAs, caution may be warranted for reports of 100+ novel miRNAs in a species (Axtell and Meyers, 2018). Further, all these flax miRNA identifications were carried out using an older scaffold-based genome (Wang et al., 2012). Recently, a chromosome-scale genome assembly has been released for flax (You et al., 2018).

To comprehensively annotate miRNA precursors in the newest flax genome, sRNA libraries were sequenced from leaves, stem, and root tissues. In addition, to identify heat responsive miRNAs in flax, sRNA transcriptomes from reproductive bud stages (Supplementary Figure 1A) at three time points were sequenced (see methods for details). In total, we generated 24 sRNA libraries, and clean reads (after removing rRNA/tRNA sequences) were used for miRNA prediction (Supplementary Table 1). To accurately identify miRNA precursors, we used ShortStack and miR-PREFeR, some of the best-performing miRNA annotation tools in terms of precision and accuracy (Johnson et al., 2016; Lei and Sun, 2014). We identified 173 candidate miRNA precursors yielding 106 mature miRNAs (Figure 1, outer circle; Supplementary Table 2). Among them, 131 precursors produced known miRNAs while 42 precursors generated novel miRNAs which contained at least 5 variances (mismatch plus unaligned nucleotides) with Viridiplantae miRNAs derived from miRBase version 22. We identified miRNAs from 31 known miRNA families, among them, the families having the greatest numbers of precursors encoded into the genome were miR167 (14), miR2118/482 (12), miR156 (11), miR166 (9), and miR160 (9) (Supplementary Figure 1B, Supplementary Table 2).

**Figure 1.**
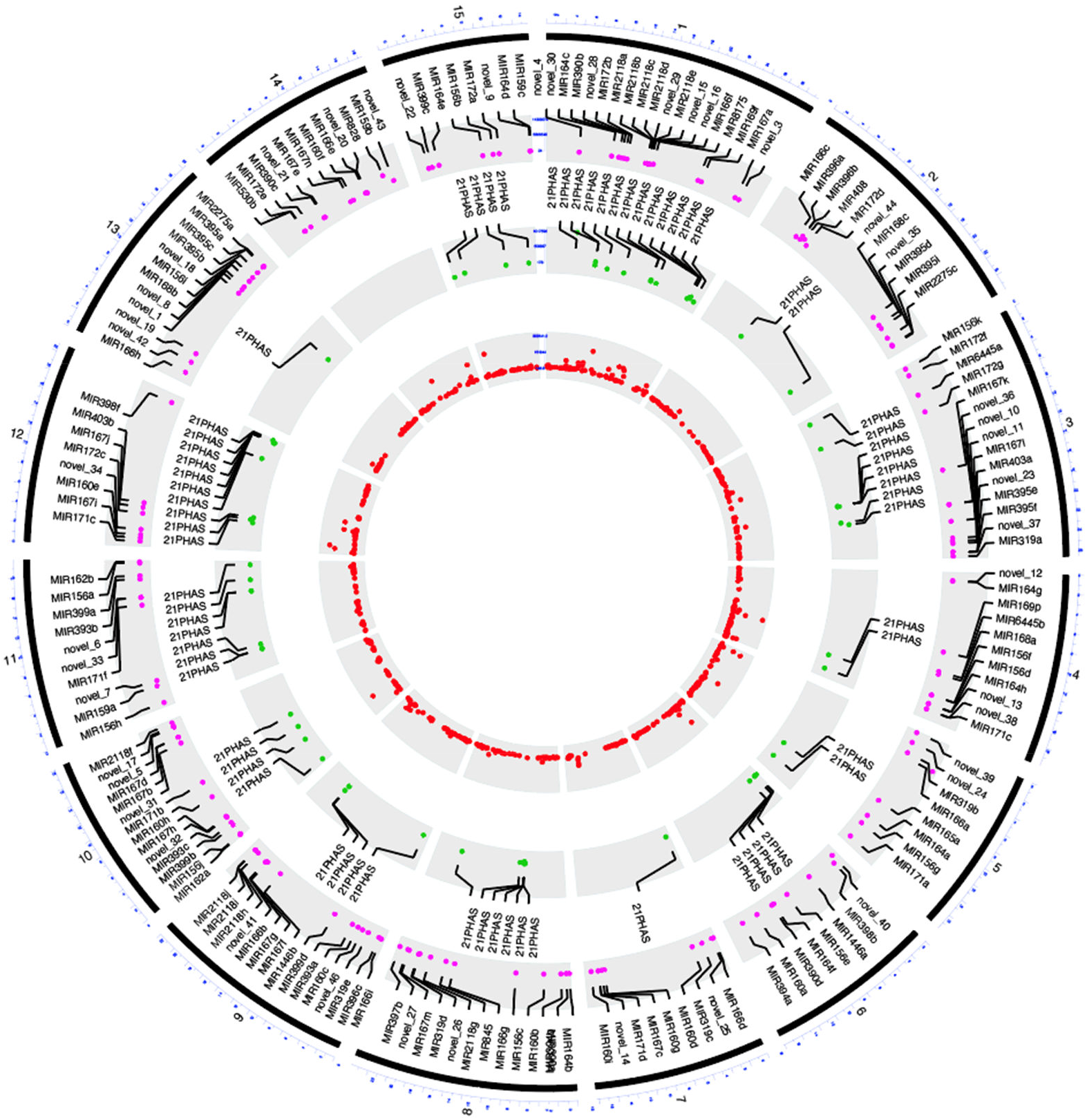
Genome-wide distribution of *MIR* genes and *PHAS* loci in flax Chromosomal position of 173 *MIR* genes (outer circle, pink dots), 68 21-*PHAS* loci (middle circle, green dots) and 662 24-*PHAS*/24-*PHAS*-like loci (inner cycle, red dots). The *MIR* genes and 21-*PHAS* loci were annotated in each chromosome. The outermost cycle represents chromosomes from 1 to 15 as numbered.

### Differentially accumulating miRNA precursors in flax tissues and heat treatments

miRNA species (miRNA-miRNA* pairs) are produced from precise cleavage of stem-loop RNA precursors, called miRNA precursors. The miRNA species accumulate differentially to perform their roles in developmental processes in vegetative and reproductive tissues. To assess miRNA precursor abundance, we summed the abundance of sRNAs that mapped to the region of the hairpin and this was used also for subsequent differential expression analyses. Hereafter, we refer to this as the ‘miRNA locus abundance’. Here, we characterized the abundance of miRNA loci in different tissue stages and summarized the results (Table 1). In total, we found 140 miRNA loci varying in abundance between vegetative and reproductive tissues; among them, 89 were enriched in vegetative tissues (Figure 2A, Supplementary Table 3) while 51 were enriched in reproductive tissues (Figure 2B, Supplementary Table 3). Set I of the vegetative tissue-enriched group, represents miRNA loci more abundant in both leaves and root, and includes members of miRNA families miR160, miR156, miR166, miR393, miR395, miR845, miR482, and several novel miRNAs (22, 26, and 46). Set II of this group is enriched mostly in root tissues and composed of members of miR164, miR156, miR395, miR159, miR399, miR394 and several novel miRNAs (20, 38, and 13). Set III, mostly enriched in both stem and root tissues, is comprised of members of miR8175, miR396, miR395, miR169, miR172, miR171, miR159 and novel miRNAs (15, 16, 25,17, 10, 14, 11, 36, 3). Set IV of this group is enriched mostly in leaves and composed of members related to miR167, miR171, miR166, miR162, miR172, miR396, miR530, miR396, miR6445, miR168, miR397, miR408, miR398 and several novel miRNAs (9, 31, 39, and 12). We divided the group of miRNA loci that are enriched more in reproductive tissues into three sets. Set I, mostly enriched in all three reproductive tissues along with stem and root, includes members of miR156, miR319, and miR2118/482. Set II of this group, mostly enriched in reproductive tissues, includes members of miR171, miR390, miR1446, miR2118/482, miR2275, miR169 and novel miRNAs (30, 24, and 7). The last Set (III) of this group were enriched in reproductive tissues and leaves, and consists of members of miR160, miR393, miR164, miR403, miR171, miR167 and novel miRNAs (5, 18, 33, 23, and 43).

**Table 1.** Differentially accumulated miRNA precursors in different tissues of flax.

|  | Leaves | Stem | Root | B_2_4 | B_5_7 | B_8_10 |
| --- | --- | --- | --- | --- | --- | --- |
| **Leaves** |  | 30D, 18U | 35D, 46U | 28D, 40U | 33D, 31U | 28D, 27U |
| **Stem** |  |  | 11D, 32U | 16D, 36U | 16D, 25U | 21D, 25U |
| **Root** |  |  |  | 45D, 42U | 51D, 39U | 53D, 31U |
| **B_2_4** |  |  |  |  | 5 D, 0U | 14D, 4U |
| **B_5_7** |  |  |  |  |  | 2D, 2U |

**Figure 2.**
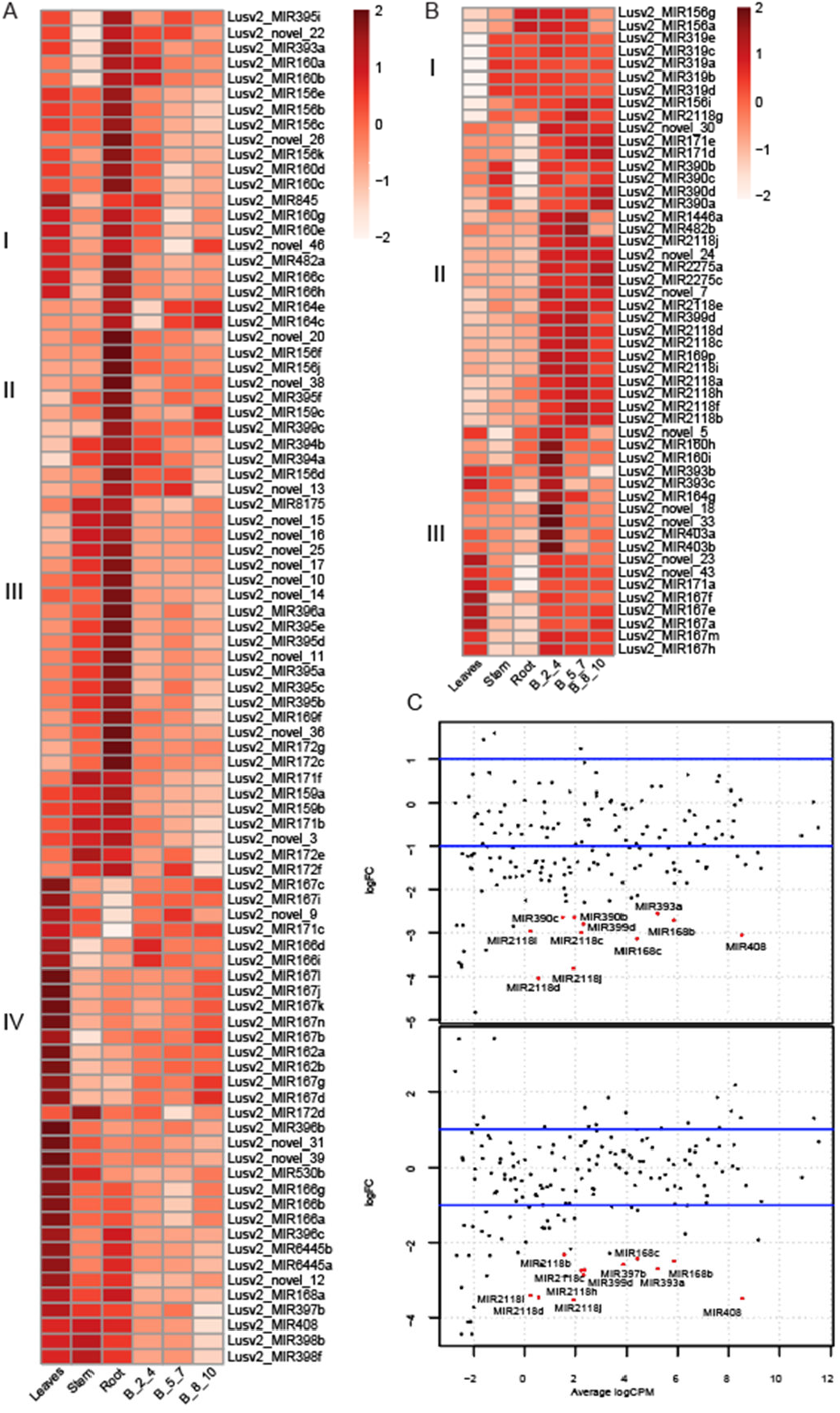
Differentially accumulated *MIR* genes in different tissues type and heat-treatments A: Heatmap showing differentially expressed *MIR* genes enriched in vegetative tissues with four sets of *MIR* genes, set I: enriched in leaves and root, set II: enriched in roots, set III: enriched in stem and roots, set IV: enriched in leaves. B. Similar to A except enriched in reproductive tissues, and with three sets of MIR genes, set I: enriched in all reproductive tissues and stem/roots, set II: enriched in reproductive tissues, and set III: enriched in reproductive tissues and leaves. The key at right shows z-score. B_2_4 indicates bud stage 1 with 2 to 4 mm in length, B_5_7 indicates bud stage 2 with 5 to 7 mm in length, and B_8_10 indicates bud stage 3 with 8 to 10 mm in length. C. Smear plot showing differentially expressed miRNAs in reproductive bud stages; upper track: bud stage 1 before (B_2_4) and after 14 days (B_2_4_F) of heat treatment, and in lower tack: bud stage 2 before (B_5_7) and after 14 days (B_5_7_F) of heat treatment.

Heat treatments applied to reproductive-stage tissues resulted in differential abundances of a total of 18 miRNA loci (Figure 2C, Supplementary Figure 2A, Supplementary Table 4). 3These were mostly downregulated due to heat stress except one: miR8175 (Supplementary Figure 2B). The downregulated miRNAs under heat stress are members of the families of miR2118, miR390, miR168, and miR393a, miR399d, miR397b, miR408, novel_7. Compared to other bud stages, bud stage 2 is the most sensitive to heat stress, and 11 miRNA loci were differentially accumulated after 7 days of heat treatment. There were no differentially accumulated miRNA loci identified between two heat treatments in all stages, but with an increased number of heat stress days, from 7 to 14, more miRNA loci were differentially expressed (Table 2). Mostly, members of miR2118/482 are enriched in reproductive tissues and they initiate the production of 21-nt reproductive phasiRNAs (Pokhrel et al., 2021a).

**Table 2.** Differentially accumulated miRNA precursors in flax due to heat treatments.

|  | B_2_4 | B_5_7 | B_8_10 | B_2_4_S | B_5_7_S | B_8_10_S | B_2_4_F | B_5_7_F | B_8_10_F |
| --- | --- | --- | --- | --- | --- | --- | --- | --- | --- |
| **B_2_4** |  |  |  | 1D, 0U |  |  | 11D, 0U |  |  |
| **B_5_7** |  |  |  |  | 10D, 1U |  |  | 12D, 0U |  |
| **B_8_10** |  |  |  |  |  | 1D, 0U |  |  | 0D, 0U |
| **B_2_4_S** |  |  |  |  |  |  | 0D, 0U |  |  |
| **B_5_7_S** |  |  |  |  |  |  |  | 0D, 0U |  |
| **B_8_10_S** |  |  |  |  |  |  |  |  | 0D, 0U |

### Genome-wide identification of phasiRNAs in flax

Plant genomes encode loci that produce secondary siRNAs that are generated from a miRNA cleavage site within a polyadenylated RNA; these are generated in a phased pattern and are called *PHAS* loci. These loci can be derived from both protein-coding (mRNA) and noncoding (long, non-coding RNA or lncRNA) transcripts. We used small RNA libraries from both vegetative and reproductive tissues to identify *PHAS* loci in flax. We identified 68 21-*PHAS* loci (i.e. loci generating 21-nt phasiRNAs) from both coding and non-coding regions (Figure 1, middle circle; Table 3; Supplementary Table 5). Of 68 loci, 51 were enriched in reproductive tissues (Figure 3A) while 17 loci were enriched in vegetative tissues (Supplementary Figure 3A). We identified three loci in flax of the most conserved phasiRNA pathway in land plants (Xia et al., 2015) miR390-*TAS3* (example locus in Figure 3B). We found two non-coding *TAS-like* (*TASL*) loci enriched in leaves and reproductive tissues triggered by miR167h (example locus in Figure 3C, miR167 variants in Figure 3D) and another 12 non-coding loci enriched in reproductive tissues with unknown triggers (Figure 3A). We found triggers of at least 30 21-*PHAS* loci, including miR167 as a trigger for a locus encoding a metal tolerance protein, miR168 for *AGO1*, miR393 for *AFB2*, miR6445 for *NAC*, miR408 for a laccase-encoding gene, and miR2118/482 for non-coding reproductive 21-*PHAS* loci that were recently described (Pokhrel et al., 2021a). The other gene families for which triggers were not identified included genes encoding BHLH, SCD, RING-type, Zf-C3H4 and BZIP domain-containing proteins, a eukaryotic translation initiation factor, and an E3 ubiquitin protein ligase (Table 3).

**Table 3.** Summary of 21-*PHAS* Loci and their triggers identified in flax.

| Trigger miRNA | Locus type, name, or encoded protein | Number of loci |
| --- | --- | --- |
| Lusv2_MIR167h/j | Metal tolerance protein | 3 |
| Lusv2_MIR167h | Non-coding | 2 |
| Lusv2_MIR168a | AGO1 | 2 |
| Lusv2_MIR2118/482 | Non-coding | 13 |
| Lusv2_MIR390 | *TAS3* | 3 |
| Lusv2_MIR393b | *AFB2* | 2 |
| Lusv2_MIR408 | Laccase | 2 |
| Lusv2_MIR6445a | *NAC* | 3 |
| N.D. | Non-coding | 13 |
| N.D. | Other protein-coding genes | 25 |

**Figure 3.**
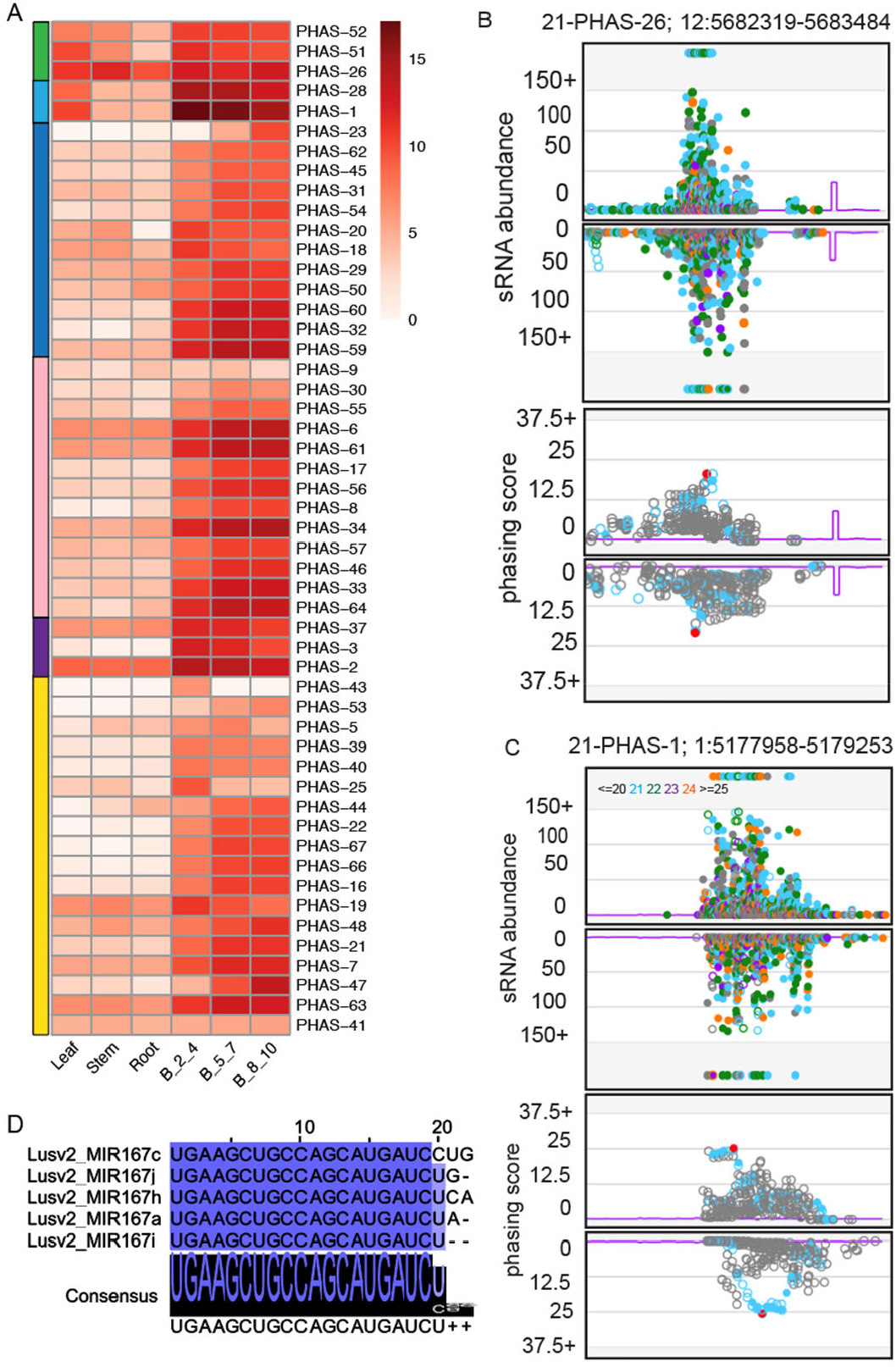
Characterization of reproductive 21-nt phasiRNAs in flax. A. Expression of reproductive-enriched multi-miRNA triggered 21-nt reproductive phasiRNAs in different tissue types. Left color-coded panel indicates different trigger miRNA and their phasiRNA; green: miR390 triggered *TAS3* loci, light blue: miR167 triggered TASL loci, dark blue: non-coding loci with unknown trigger, non-coding loci with miR2118/482 trigger, purple: miR167 derived metal proteins related loci, and yellow: all other protein coding loci with unknown triggers. The key at right indicates the abundance in unit of log2(RP20M). B_2_4, B_5_7, and B_8_10 indicate buds stage 1, 2 and 3 respectively. B. Upper track: Abundance (RP10M) of sRNAs in both strands of an example TAS3 locus padded with 250 base pairs, each side. Name and coordinates of the *PHAS* locus are indicated in top. sRNA sizes are denoted by colors, as indicated at top. Below track: Phasing score of the same locus; the red dot indicates the highest phased sRNA position. C. Similar to B for an example TASL locus as indicated in the top. D. Alignment of members of miR167 in rose. The intensity of blue color denotes the conservation in nucleotide level. The consensus sequence of the alignment is shown with a sequence logo.

Recently, 24-nt phasiRNAs were reported in several eudicots (Xia et al., 2019; Pokhrel et al., 2021b), thus, we looked for the conservation of this pathway in flax. We identified four 24-*PHAS* loci triggered by miR2275 and an additional 658 24-*PHAS*-like loci with unknown triggers (Figure 1, inner circle; Supplemental Figure 3B; Supplementary Table 6). These 24-*PHAS*-like loci passed the software’s filtering criteria for *PHAS* loci and they were mostly derived from non-repetitive regions of the genome; however, when we examined a subset in our genome browser (Nakano et al., 2020), we found these loci display poor phasing, with phasing scores ranging from 15 to 20 .Due to their poor phasing and unknown triggering mechanism, It is more likely that these 24-nt sRNAs could be hc-siRNAs that are abundant at specific stage of reproductive development, misidentified as 24-nt phasiRNAs as previously described (Polydore et al., 2018). However, subset of these loci may be derived from 12-nt phasing similar to the 24-nt phasiRNAs of Solanaceous species (Xia et al., 2019).

### Differentially expressed phasiRNAs

Since the two classes of phasiRNAs enriched in anthers of angiosperms demonstrate short periods of peak abundance (Xia et al., 2019; Pokhrel et al., 2021a, 2021b), we examined the differential accumulation of these reproductive phasiRNAs. We found among 68 21-*PHAS* loci present in flax that 40 loci were differentially expressed, with most (37) being downregulated in heat-treated samples (Supplemental Table 7). Among the differentially expressed loci, 37 loci were enriched in reproductive tissues, 12 of which are 21-*PHAS* loci targeted by miR2118/482, three encode metal tolerance proteins (triggered by miR167h/f), nine are non-coding loci with unknown triggers, and remaining 13 loci are from other protein-coding genes (Figure 4A, Supplemental Table 7). Both stages 1 and 2 had similar numbers of 21-*PHAS* differentially expressed during 7 and 14 days of heat treatments, however for stage 3, the 14-day heat treatment increased the number of differentially expressed number of 21-*PHAS* loci (Table 4).

**Figure 4.**
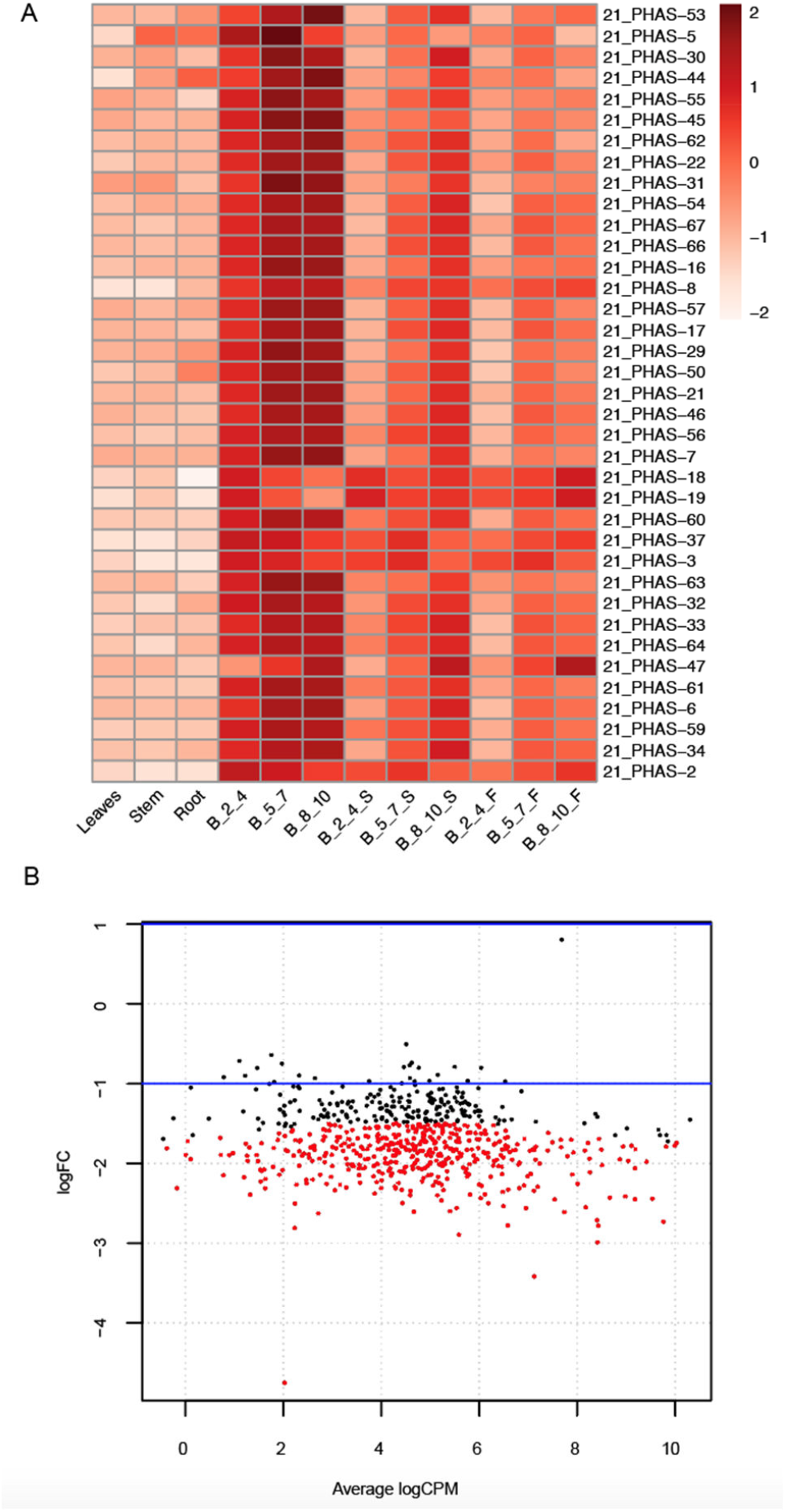
Profiling of differentially accumulation of reproductive 21 and 24-nt phasiRNAs in flax. A. Differential accumulation of 21-nt reproductive phasiRNAs in reproductive stages after heat treatments. B_2_4, B_5_7, and B_8_10 indicate bud stage 1, 2 and 3 respectively. S and F indicates seven and fourteen days of heat treatment. The key at right indicates z-score. B. Smear plot showing differentially expressed 24-nt phasiRNAs in bud stage 1 before (B_2_4) and after 14 days (B_2_4_F) of heat treatment.

**Table 4.**
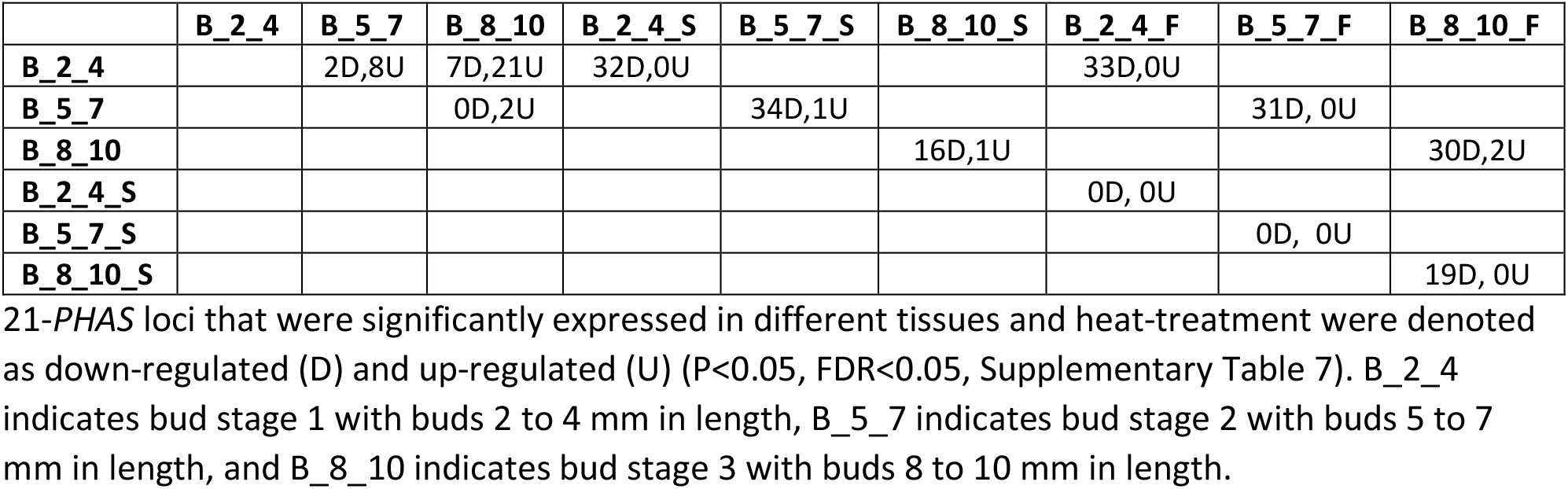
Differentially expressed 21-*PHAS* loci in flax due to heat-treatments.

|  | B_2_4 | B_5_7 | B_8_10 | B_2_4_S | B_5_7_S | B_8_10_S | B_2_4_F | B_5_7_F | B_8_10_F |
| --- | --- | --- | --- | --- | --- | --- | --- | --- | --- |
| B_2_4 |  | 2D,8U | 7D,21U | 32D,0U |  |  | 33D,0U |  |  |
| B_5_7 |  |  | 0D,2U |  | 34D,1U |  |  | 31D, 0U |  |
| B_8_10 |  |  |  |  |  | 16D,1U |  |  | 30D,2U |
| B_2_4_S |  |  |  |  |  |  | 0D, 0U |  |  |
| B_5_7_S |  |  |  |  |  |  |  | 0D, 0U |  |
| B_8_10_S |  |  |  |  |  |  |  |  | 19D, 0U |

In case of 24-*PHAS* loci, for bud stages 1 and 2, 14 days of heat treatment resulted in more downregulated 24-*PHAS* loci compared to 7 days of heat treatment, while expression of 24-*PHAS* loci in bud stage 3 didn’t change due to either of the heat-treatments (Table 5). Each of the four reproductive 24-*PHAS* loci triggered by miR2275 were down-regulated in heat-treated samples, while 450 24-*PHAS*-like loci were also down-regulated by heat, out of 658 total loci (Figure 4B, Supplemental Table 8).

**Table 5.** Differentially expressed 24-*PHAS* loci in flax due to heat-treatments.

|  | B_4_2 | B_5_7 | B_8_10 | B_2_4_S | B_5_7_S | B_8_10_S | B_2_4_F | B_5_7_F | B_8_10_F |
| --- | --- | --- | --- | --- | --- | --- | --- | --- | --- |
| B_2_4 |  |  |  | 62D,0U |  |  | 449D,0U |  |  |
| B_5_7 |  |  |  |  | 68D,0U |  |  | 143D, 0U |  |
| B_8_10 |  |  |  |  |  | 1D,0U |  |  | 0D,0U |
| B_2_4_S |  |  |  |  |  |  | 0D, 0U |  |  |
| B_5_7_S |  |  |  |  |  |  |  | 0D, 0U |  |
| B_8_10_S |  |  |  |  |  |  |  |  | 0D, 0U |

## Discussion

We conducted a genome-wide annotation of miRNAs and phasiRNAs in flax. Based on updated criteria of miRNA annotation (Axtell and Meyers, 2018), we comprehensively identified 173 *MIR* genes that generated miRNAs from 31 known families and 42 novel miRNAs. As miRNAs play important regulatory roles in plant growth and reproductive development, we found that most of the miRNAs (141 out of 173) were preferentially accumulated in a tissue-specific manner. Further, we categorized differentially accumulated miRNAs based on their enrichment in vegetative and reproductive tissues. Since miRBase (v22) contains only 124 *MIR* genes for flax, most of which were identified using computational prediction based on the flax genome sequence, our current analysis would substantially improve the miRBase data for flax. We identified 68 21-*PHAS* loci that include three *TAS3* loci, two *TASL* loci, and additional loci from coding and non-coding regions. We found no *PHAS* loci related to *PPR* genes or *NB-LRR* defense genes, which is surprising as these are conserved and numerous *PHAS* loci in many eudicots. However, we identified several miR2118/482 members that initiate the production of reproductive tissue-enriched 21-nt phasiRNAs from non-coding loci in flax (Zhai et al., 2011; Pokhrel et al., 2021a).We identified several miRNAs and phasiRNA pathways in flax that are responsive to heat stress. The two members of miR390, the conserved trigger of *TAS*3, were down-regulated under to heat-stress while *TAS*3 derived phasiRNAs weren’t differentially expressed. We found no miR167 members to be differentially accumulated under to heat stress but we observed a reduction in phasiRNAs from miR167-triggered metal tolerance proteins phasing pathways were enriched in reproductive tissues and phasiRNAs were down-regulated.

In recent studies, conservation of miR2275-24-*PHAS* pathway in angiosperms were reported, including several eudicots (Xia et al., 2019; Pokhrel et al., 2021b), and the pathway is also conserved in flax, but with very few (4) 24-*PHAS* loci. Here, we identified a larger number (658) of 24-*PHAS*-like loci that are reproductive tissue-enriched but that lack obvious trigger miRNAs, similar to the Solanaceous 24-nt phasiRNAs (Xia et al., 2019). We found that all 24-*PHAS* and 450 24-*PHAS*-like loci were heat-responsive and all are down-regulated due to heat stress.

Notably, we found down-regulation of the reproductive phasiRNA triggers, miR2118/482 and miR2275 in response to heat treatments, as well as the reproductive 21- and 24-nt phasiRNAs triggered by these miRNAs respectively. In rice, mutation in the precursor of 21-*PHAS* loci resulted in temperature-sensitive genic male sterility (Fan et al., 2016; Zhou et al., 2012) while in maize, knockdown of *DCL*5, a pathway gene responsible for the production of reproductive 24-nt phasiRNAs in monocots, greatly reduced the phasiRNAs which resulted in male sterile phenotype under heat restrictive (28/22°C) temperatures (Teng et al., 2020). In this study, we subjected flax to two heat-treatments during flowering that reduced the number of boll formation and seed set as reported in (Cross et al., 2003); however the molecular mechanisms underlying these results were unknown. We hypothesize that a decrease in abundance of rate-limiting trigger miRNAs due to heat stress results in the down-regulation of reproductive phasiRNAs which ultimately produces sterility phenotypes in flax. Future studies are needed to elucidate a mechanistic understanding of the roles for miRNA-phasiRNAs in reproductive development under heat stress.

## Methods

### Heat stress treatment and plant materials

Flax cultivar ‘Stormont cirrus’ was grown in a walk-in Conviron growth chamber with 16/8 hour light/dark cycle with 25/20 °C temperatures, respectively. From one month-old seedlings, leaves, stem, and root tissues were harvested and immediately frozen in liquid nitrogen. After two weeks of initiation of flowering, two groups of seedlings, each containing at least nine plants, were transferred to a new growth chamber where during the day (16 h) cycle, temperatures would increase at ∼3°C /h from 20 °C to 40 °C, over a 7 h period, hold at 40 °C for 1 h, and decrease at the same rate for 7 h. The heat treatment was continued for fourteen days and samples of three bud stages (Supplementary Figure 1A) were harvested before initiation of heat stress, and after seven and fourteen days during peak heat stress (40 °C), from two groups as two biological replicates, and frozen immediately in liquid nitrogen.

### RNA extraction, libraries preparation and sequencing

Total RNA from all flax vegetative and reproductive tissues was extracted using the PureLink Plant RNA Reagent (ThermoFisher Scientific, cat. #12322012) following the manufacturer’s instructions with minor modifications. RNA quality was determined by running denaturing gels and quantity was assessed using Qubit. First, sRNAs (20 to 30 nt) were size-selected in a 15% polyacrylamide/urea gel and used for sRNA library preparation. An input amount 5 μg of total RNA was used for library construction. The sRNA libraries were constructed following manufacturer’s instruction using the NEBNext Small RNA Library Preparation set for Illumina (NEB, cat # E7300).

### Data analysis and visualization

Pre-processing and filtering of sRNA libraries were conducted as previously described (Pokhrel et al., 2021a). Two miRNA identification tools: ShortStack and miR-PREFeR were used (Lei and Sun, 2014; Johnson et al., 2016) for the genome-wide MIR genes identification. To increase the number of miRNA detection, in ShortStack, two libraries were used at a time using default parameters except ---mismatch 0 and --mincov 10. As ShortStack is the tool with highest precision and accuracy, miRNA loci predicted using miR-PREFeR were again re-evaluated with ShortStack and only the loci with passed the criteria were included in the final set. The miRNA loci from two tools were merged using BEDTools (Quinlan and Hall, 2010).

*PHAS* loci were also identified using two tools, *PHASIS* (Kakrana et al., 2017) with a p-value cut-off of 0.001 and number of phase cycle (k) =7 and ShortStack (Johnson et al., 2016) with default parameters except –mismatch 0 and –mincov 10. BEDTools was used to merge *PHAS* loci from two different softwares (Quinlan and Hall, 2010). 24-*PHAS* loci with unknown triggers were annotated as 24-*PHAS* –like loci. Target prediction was carried out by sPARTA (Kakrana et al., 2014) with a target cut off score of ≤5. The sRNA abundance and phasing score were viewed at a customized browser (Nakano et al., 2020). The 250 base pair padded sequence of *PHAS* loci were annotated de novo by BLASTX (Altschul et al., 1997) against UniRef90. Differential analysis (*P*<0.05, FDR<0.05) of sRNAs were conducted in R using the ‘edgeR’ package (Robinson et al., 2010). Circular plots were constructed using OmicCircos (Hu et al., 2014) for the chromosomal distributions of miRNAs and phasiRNAs generating loci. pheatmap (https://rdrr.io/cran/pheatmap/) was used to draw heatmaps in R.

## Acknowledgements

We thank members of the Meyers lab for helpful discussions, and Joanna Friesner for assistance with editing. We thank Mayumi Nakano for assistance with data handling. This work was supported by resources from the Donald Danforth Plant Science Center and the University of Missouri – Columbia plus the William H. Danforth Plant Science Fellow Award (to S.P.).

## Author Contributions

B.C.M. and S.P. designed the research; S.P. performed the experiments and analyzed the data; S.P. and B.C.M wrote and revised the manuscript.

## Supporting Information

**Supplemental Figure 1. Reproductive buds staging and miRNA precursors in flax**

A. Bud stages used for sequencing in flax, B_2_4: 2-4 mm buds, B_5_7: 5-7mm buds, and B_8_10: 8-10mm buds. B. Number of precursors encoded into the genome of flax for each miRNA families. miRNA family with more than 3 precursors were included.

**Supplemental Figure 2. Differential accumulation of miRNAs due to heat stress in flax**.

A. Differential accumulation of miRNAs in reproductive buds after heat stress. The key at right indicates z-score. B_2_4: 2-4 mm buds, B_5_7: 5-7mm buds, and B_8_10: 8-10mm buds. S and F indicates seven and fourteen days of heat treatment.

B. Smear plot showing differentially expressed miRNAs in bud stage 2 (before B_5_7) and after 7 days (B_5_7_S) of heat treatment.

**Supplemental Figure 3. Accumulation of vegetative-enriched 21-nt phasiRNAs and all 24-nt phasiRNA in flax**

A. Accumulation pattern of vegetative-enriched 21-nt phasiRNAs in different tissues. The key at right indicates the abundance in units of log2(RP20M). B_2_4: 2-4 mm buds, B_5_7: 5-7mm buds, and B_8_10: 8-10mm buds.

B. Accumulation of 24-nt reproductive phasiRNAs derived from 24-PHAS/24-PHAS-like loci in different tissues. The key at right indicates the abundance in units of log2(RP20M). B_2_4: 2-4 mm buds, B_5_7: 5-7mm buds, and B_8_10: 8-10mm buds.

**Supplemental Figure 4. 24-nt phasiRNAs differentially expressed after heat-stress treatment in flax**.

Differential accumulation of 24-nt phasiRNAs from 454 loci in reproductive buds after heat stress. The key at right indicates z-score. B_2_4: 2-4 mm buds, B_5_7: 5-7mm buds, and B_8_10: 8-10mm buds. S and F indicates seven and fourteen days of heat treatment.

## References

Altschul, S.F., Madden, T.L., Schäffer, A.A., Zhang, J., Zhang, Z., Miller, W., and Lipman, D.J. (1997). Gapped BLAST and PSI-BLAST: A new generation of protein database search programs. Nucleic Acids Res. 25: 3389–3402.

Axtell, M.J. and Meyers, B.C. (2018). Revisiting criteria for plant microRNA annotation in the Era of big data. Plant Cell 30: 272–284.

Barvkar, V.T., Pardeshi, V.C., Kale, S.M., Qiu, S., Rollins, M., Datla, R., Gupta, V.S., and Kadoo, N.Y. (2013). Genome-wide identification and characterization of microRNA genes and their targets in flax (Linum usitatissimum): Characterization of flax miRNA genes. Planta 237: 1149–1161.

Cross, R.H., McKay, S.A.B., McHughen, A.G., and Bonham-Smith, P.C. (2003). Heat-stress effects on reproduction and seed set in Linum usitatissimum L. (flax). Plant, Cell Environ. 26: 1013–1020.

Fan, Y. et al. (2016). PMS1T, producing phased small-interfering RNAs, regulates photoperiod-sensitive male sterility in rice. Proc. Natl. Acad. Sci. 113: 15144–15149.

Fei, Q., Xia, R., and Meyers, B.C. (2013). Phased, secondary, small interfering RNAs in posttranscriptional regulatory networks. Plant Cell 25: 2400–2415.

Giacomelli, J.I., Weigel, D., Chan, R.L., and Manavella, P.A. (2012). Role of recently evolved miRNA regulation of sunflower HaWRKY6 in response to temperature damage. New Phytol. 195: 766–773.

Guan, Q., Lu, X., Zeng, H., Zhang, Y., and Zhu, J. (2013). Heat stress induction of miR398 triggers a regulatory loop that is critical for thermotolerance in Arabidopsis. Plant J. 74: 840–851.

Hasanuzzaman, M., Nahar, K., Alam, M.M., Roychowdhury, R., and Fujita, M. (2013). Physiological, biochemical, and molecular mechanisms of heat stress tolerance in plants. Int. J. Mol. Sci. 14: 9643–9684.

He, J., Jiang, Z., Gao, L., You, C., Ma, X., Wang, X., Xu, X., Mo, B., Chen, X., and Liu, L. (2019). Genome-wide transcript and small RNA profiling reveals transcriptomic responses to heat stress. Plant Physiol. 181: 609–629.

Hedhly, A., Hormaza, J.I., and Herrero, M. (2009). Global warming and sexual plant reproduction. Trends Plant Sci. 14: 30–36.

Hu, Y., Yan, C., Hsu, C.H., Chen, Q.R., Niu, K., Komatsoulis, G.A., and Meerzaman, D. (2014). Omiccircos: A simple-to-use R package for the circular visualization of multidimensional Omics data. Cancer Inform. 13: 13–20.

Johnson, N.R., Yeoh, J.M., Coruh, C., and Axtell, M.J. (2016). Improved placement of multi-mapping small RNAs. G3 Genes, Genomes, Genet. 6: 2103–2111.

Kakrana, A., Hammond, R., Patel, P., Nakano, M., and Meyers, B.C. (2014). SPARTA: A parallelized pipeline for integrated analysis of plant miRNA and cleaved mRNA data sets, including new miRNA target-identification software. Nucleic Acids Res. 42: 1–13.

Kakrana, A., Li, P., Patel, P., Hammond, R., Anand, D., Mathioni, S.M., and Meyers, B.C. (2017). PHASIS: A computational suite for de novo discovery and characterization of phased, siRNA-generating loci and their miRNA triggers. bioRxiv.

Keller, M., Schleiff, E., and Simm, S. (2020). miRNAs involved in transcriptome remodeling during pollen development and heat stress response in Solanum lycopersicum. Sci. Rep. 10: 10694.

Lei, J. and Sun, Y. (2014). miR-PREFeR: an accurate, fast and easy-to-use plant miRNA prediction tool using small RNA-Seq data. Bioinformatics 30: 2837–2839.

Li, S., Liu, J., Liu, Z., Li, X., Wu, F., and He, Y. (2014). HEAT-INDUCED TAS1 TARGET1 mediates thermotolerance via heat stress transcription factor A1a-directed pathways in arabidopsis. Plant Cell 26: 1764–1780.

Lin, J.S., Kuo, C.C., Yang, I.C., Tsai, W.A., Shen, Y.H., Lin, C.C., Liang, Y.C., Li, Y.C., Kuo, Y.W., King, Y.C., Lai, H.M., and Jeng, S.T. (2018). MicroRNA160 modulates plant development and heat shock protein gene expression to mediate heat tolerance in Arabidopsis. Front. Plant Sci. 9: 68.

Melnikova, N. V. et al. (2016). Identification, expression analysis, and target prediction of flax genotroph MicroRNAs under normal and nutrient stress conditions. Front. Plant Sci. 7: 1–12.

Nakano, M., McCormick, K., Demirci, C., Demirci, F., Gurazada, S.G.R., Ramachandruni, D., Dusia, A., Rothhaupt, J.A., and Meyers, B.C. (2020). Next-generation sequence databases: RNA and genomic informatics resources for plants. Plant Physiol. 182: 136–146.

Pokharel, M., Chiluwal, A., Stamm, M., Min, D., Rhodes, D., and Jagadish, S.V.K. (2020). High night-time temperature during flowering and pod filling affects flower opening, yield and seed fatty acid composition in canola. J. Agron. Crop Sci. 206: 579–596.

Pokhrel, S., Huang, K., Bélanger, S., Zhan, J., Caplan, J.L., Kramer, E.M., and Meyers, B.C. (2021a). Pre-meiotic 21-nucleotide reproductive phasiRNAs emerged in seed plants and diversified in flowering plants. Nat. Commun. 12.

Pokhrel, S., Huang, K., and Meyers, B.C. (2021b). Conserved and non-conserved triggers of 24-nucleotide reproductive phasiRNAs in eudicots. Plant J.: 1–14.

Polydore, S., Lunardon, A., and Axtell, M.J. (2018). Several phased siRNA annotation methods can frequently misidentify 24 nucleotide siRNA-dominated PHAS loci. Plant Direct 2.

Quinlan, A.R. and Hall, I.M. (2010). BEDTools: A flexible suite of utilities for comparing genomic features. Bioinformatics 26: 841–842.

Robinson, M.D., McCarthy, D.J., and Smyth, G.K. (2010). edgeR: a Bioconductor package for differential expression analysis of digital gene expression data. Bioinformatics 26: 139–140.

Stief, A., Altmann, S., Hoffmann, K., Pant, B.D., Scheible, W.R., and Bäurle, I. (2014). Arabidopsis miR156 regulates tolerance to recurring environmental stress through SPL transcription factors. Plant Cell 26: 1792–1807.

Teng, C., Zhang, H., Hammond, R., Huang, K., Meyers, B.C., and Walbot, V. (2020). Dicer-like 5 deficiency confers temperature-sensitive male sterility in maize. Nat. Commun. 11: 2912.

Wang, D., Heckathorn, S.A., Mainali, K., and Tripathee, R. (2016). Timing effects of heat-stress on plant ecophysiological characteristics and growth. Front. Plant Sci. 7: 1629.

Wang, Z. et al. (2012). The genome of flax (Linum usitatissimum) assembled de novo from short shotgun sequence reads. Plant J. 72: 461–473.

Xia, R., Chen, C., Pokhrel, S., Ma, W., Huang, K., Patel, P., Wang, F., Xu, J., Liu, Z., Li, J., and Meyers, B.C. (2019). 24-nt reproductive phasiRNAs are broadly present in angiosperms. Nat. Commun. 10: 627.

Xia, R., Xu, J., Arikit, S., and Meyers, B.C. (2015). Extensive families of miRNAs and PHAS loci in Norway spruce demonstrate the origins of complex phasiRNA networks in seed plants. Mol. Biol. Evol. 32: 2905–2918.

You, F.M., Xiao, J., Li, P., Yao, Z., Jia, G., He, L., Zhu, T., Luo, M.C., Wang, X., Deyholos, M.K., and Cloutier, S. (2018). Chromosome-scale pseudomolecules refined by optical, physical and genetic maps in flax. Plant J. 95: 371–384.

Yu, Y., Wu, G., Yuan, H., Cheng, L., Zhao, D., Huang, W., Zhang, S., Zhang, L., Chen, H., Zhang, J., and Guan, F. (2016). Identification and characterization of miRNAs and targets in flax (Linum usitatissimum) under saline, alkaline, and saline-alkaline stresses. BMC Plant Biol. 16.

Zhai, J. et al. (2011). MicroRNAs as master regulators of the plant NB-LRR defense gene family via the production of phased, trans-acting siRNAs. Genes Dev. 25: 2540–2553.

Zhang, S.S., Yang, H., Ding, L., Song, Z.T., Ma, H., Chang, F., and Liu, J.X. (2017). Tissue-specific transcriptomics reveals an important role of the unfolded protein response in maintaining fertility upon heat stress in Arabidopsis. Plant Cell 29: 1007–1023.

Zhao, C. et al. (2017). Temperature increase reduces global yields of major crops in four independent estimates. Proc. Natl. Acad. Sci. U. S. A. 114: 9326–9331.

Zhou, H. et al. (2012). Photoperiod- and thermo-sensitive genic male sterility in rice are caused by a point mutation in a novel noncoding RNA that produces a small RNA. Cell Res. 22: 649–660.

