## Supplemental Figure for "Characterization of Heat Responsive microRNAs and Phased Small Interfering RNAs in Reproductive Development of Flax"

**Supplemental Figure 1. Reproductive buds staging and miRNA precursors in flax**

A. Bud stages used for sequencing in flax, B_2_4: 2-4 mm buds, B_5_7: 5-7mm buds, and B_8_10: 8-10mm buds. B. Number of precursors encoded into the genome of flax for each miRNA families. miRNA family with more than 3 precursors were included.

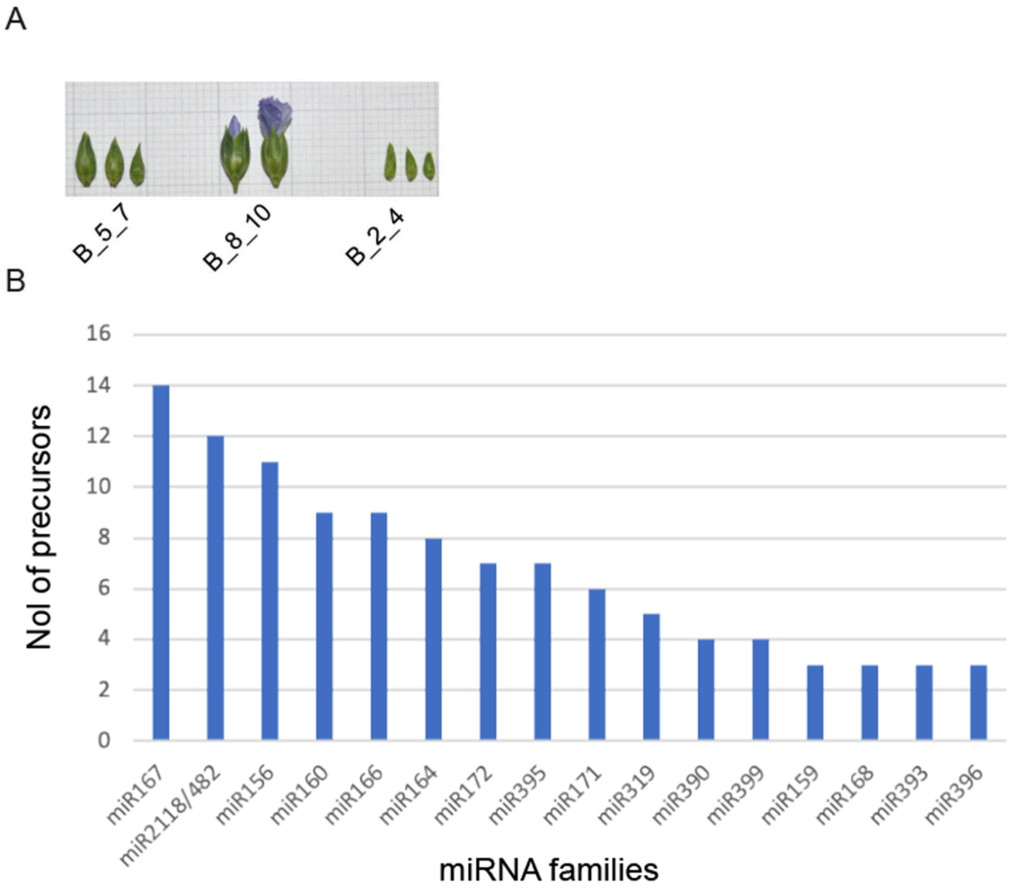

**Supplemental Figure 2. Differential accumulation of miRNAs due to heat stress in flax.**

A. Differential accumulation of miRNAs in reproductive buds after heat stress. The key at right indicates z-score. B_2_4: 2-4 mm buds, B_5_7: 5-7mm buds, and B_8_10: 8-10mm buds. S and F indicates seven and fourteen days of heat treatment.

B. Smear plot showing differentially expressed miRNAs in bud stage 2 before (B_5_7) and after 7 days (B_5_7_S) of heat treatment.

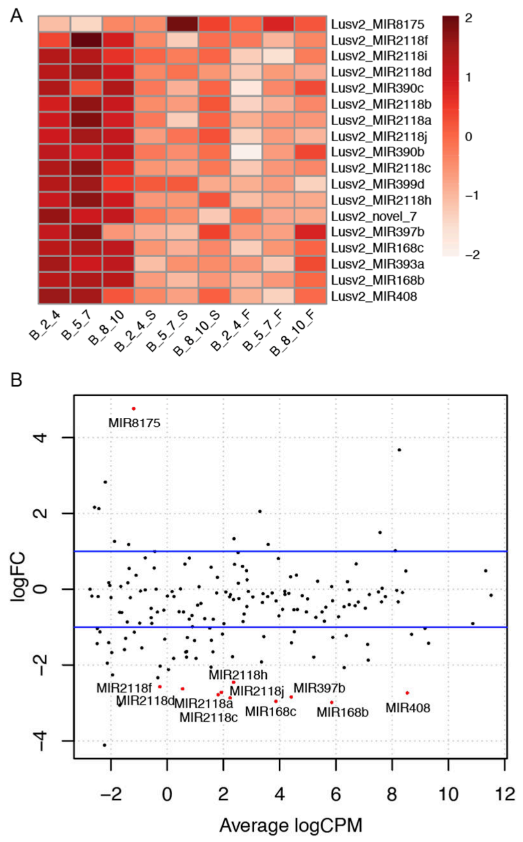

**Supplemental Figure 3. Accumulation of vegetative-enriched 21-nt phasiRNAs and all 24-nt phasiRNA in flax**

A. Accumulation pattern of vegetative-enriched 21-nt phasiRNAs in different tissues. The key at right indicates the abundance in units of log2(RP20M). B_2_4: 2-4 mm buds, B_5_7: 5-7mm buds, and B_8_10: 8-10mm buds.

B. Accumulation of 24-nt reproductive phasiRNAs derived from 24-PHAS/24- PHAS-like loci in different tissues. The key at right indicates the abundance in units of log2(RP20M). B_2_4: 2-4 mm buds, B_5_7: 5-7mm buds, and B_8_10: 8-10mm buds.

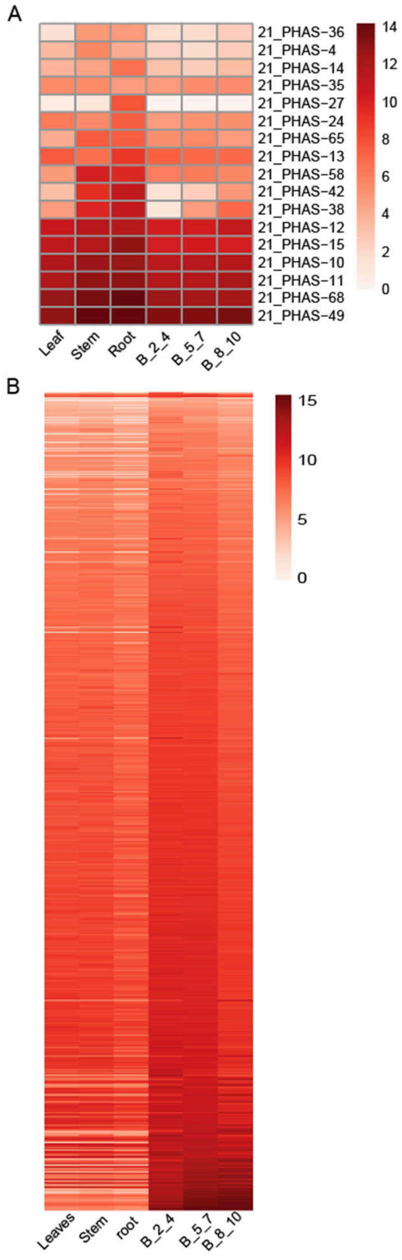

**Supplemental Figure 4. 24-nt phasiRNAs differentially expressed after heat-stress treatment in flax.**

Differential accumulation of 24-nt phasiRNAs from 454 loci in reproductive buds after heat stress. The key at right indicates z-score. B_2_4: 2-4 mm buds, B_5_7: 5-7mm buds, and B_8_10: 8-10mm buds. S and F indicates seven and fourteen days of heat treatment.

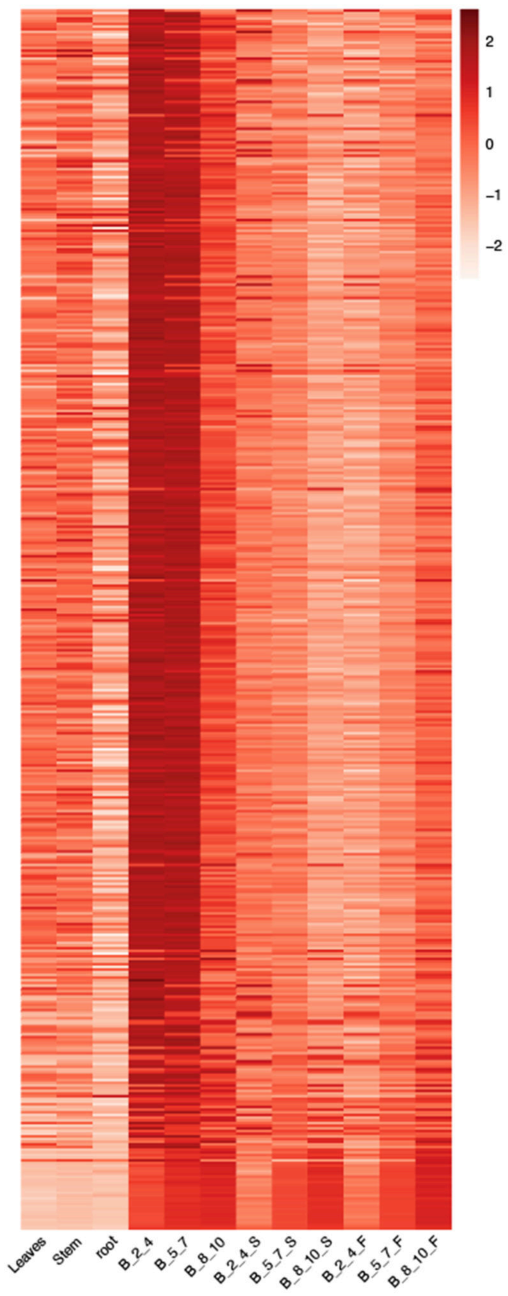
